# Influence of environmental variables, small-scale fisheries and vessel traffic on the distribution and behavior of bottlenose dolphins in a tropical lagoon

**DOI:** 10.1101/2024.02.29.582495

**Authors:** M. Brevet, S. Jaquemet, J. Wagner, JJ. Kiszka

**Author notes:** Corresponding author: Mathieu Brevet.

## Abstract

The distribution of marine predators is influenced by a variety of natural and, in some cases, anthropogenic environmental factors. In particular, the optimal foraging theory predicts that micro-habitat selection should be the result of a trade-off between prey availability, energy expenditure, and predation risk. In addition, the risk-disturbance hypothesis suggests that anthropogenic disturbance may be perceived by animals in the same way as predation risk. Habitat selection may also be locally influenced by individual behavior and physiological state (defining functional habitats): variation in their needs should affect their optimal trade-off. We tested these hypotheses in a population of bottlenose dolphins living in a tropical lagoon using a habitat modeling approach. Bottlenose dolphins were predominantly distributed within the lagoon, with a preference for the vicinity of fringing and inner reefs (with lower predation risk than the outer reef), and were located in areas of high fish productivity, consistent with optimal habitat selection. We also observed an interaction between habitat and dolphin behavior, suggesting the existence of functional habitats: foraging was more common in nearshore habitats with probable higher prey density while resting and socializing were more common further from shore. Similarly, females with calves were preferentially found in shallower waters compared to other social groups. We did not observe any effects of anthropogenic disturbance variables and therefore cannot conclude on the risk-disturbance hypothesis for this population.

## Introduction

Marine predators can be critical to the structure and function of communities and ecosystems [1]. Therefore, their conservation is crucial to maintain predator-prey interactions and nutrient transfer across ecosystem boundaries ([2], [3]). Consequently, understanding the spatial distribution of such species should be a priority to support conservation efforts.

Environmental variables are key elements in understanding the distribution and abundance of marine predators. Several eco-geographic variables are known to be relevant predictors of their distribution and abundance, in particular physiography ([4]), mesoscale oceanographic structures (eddies, upwelling, seamounts; [5], [6], [7]) and biotic conditions, including potential prey abundance (or proxies of) ([8], [9]) and predation risk [10].

The distribution of such species as a function of environmental context could be explained by theoretical frameworks such as optimal foraging theory [11] or ideal free distribution ([12]): individuals are expected to choose their habitat according to a trade-off between environmental constraints (energy expenditure, competition, predation risk) and energetic benefits (i.e. prey availability). These trade-offs have been extensively studied in terrestrial ecosystems, but are still under investigation in marine environments ([10], [13]). In such a framework, anthropogenic disturbances could also affect this trade-off, as they can be perceived in the same way as predation risk (Risk Disturbance Hypothesis; [14]). Thus, anthropogenic disturbances such as noise pollution or boat traffic could have an impact on the distribution and abundance of marine predators ([15], [16]). Moreover, given the heterogeneity of individual traits, habitat selection may vary among individuals and their activities: several functional habitats may be distinguished depending on individual needs ([12]).

We wanted to test the existence of such trade-offs and functional habitats in the Indo-Pacific bottlenose dolphin (*Tursiops aduncus*), in the tropical lagoon of Mayotte. This population is geographically and genetically isolated [17], without underlying genetic structure [18]. A recent study showed that this population is currently endangered according to the UICN Red List criteria [19]. A variety of anthropogenic impacts are suspected to be affecting this population [19], including prey depletion due to overfishing and disturbance due to increased boat traffic, dolphin-watching activities, entanglement in fishing gear [20], and habitat degradation ([21], [22]). This population is also affected by a lobomycosis-like disease outbreak, possibly related to coastal urbanization with untreated freshwater runoff [22]. Thus, this species is at relatively high risk of population decline in this region.

To effectively mitigate threats to coastal dolphin populations, it is critical to have a good understanding of their habitat preferences and the relative importance of abiotic and biotic parameters on their distribution. Thus, the identification of factors that shape their distribution and activity budgets is fundamental for conservation and management. Therefore, this study aimed to identify such factors (including anthropogenic impacts) and to understand how they influence dolphin behavior (foraging, resting, traveling, socializing) and group structure (age and sex composition). To this end, we developed generalized linear models (GLMs) to test the influence of environmental and anthropogenic factors on the distribution and behavior of Indo-Pacific bottlenose dolphins in the waters of Mayotte.

## Materials & Methods

### Study area and survey design

The Indo-Pacific bottlenose dolphin (*Tursiops aduncus*) occurs in coastal tropical and warm temperate waters from eastern Africa to the western Pacific Ocean [23]. In the waters of the Mozambique Channel island of Mayotte (SW Indian Ocean, see figure 1.A), Indo-Pacific bottlenose dolphins are distributed within the lagoon and are the most abundant dolphin species occurring around the island [24]. Small boat surveys were conducted in the coastal waters of Mayotte (45^*°*^10’E/12^*°*^50’S), including the inner waters of the lagoon, the outer slope waters of the barrier reef, and the shallow waters of the Iris Bank (figure 1.A). Surveys were conducted between 2004 and 2009 ([24], [19] for sampling details) and between 2014 and 2015. Between 2004 and 2009, samples were randomly distributed around the island to investigate the distribution, habitat preferences, and relative abundance of cetaceans (e.g. [25], [24], [18]). Between 2014 and 2015, surveys followed predetermined transects targeting coastal dolphins (primarily *Tursiops aduncus*) to homogenize sampling throughout the study area where bottlenose dolphins are most abundant ([18], [19]).

**Figure 1:**
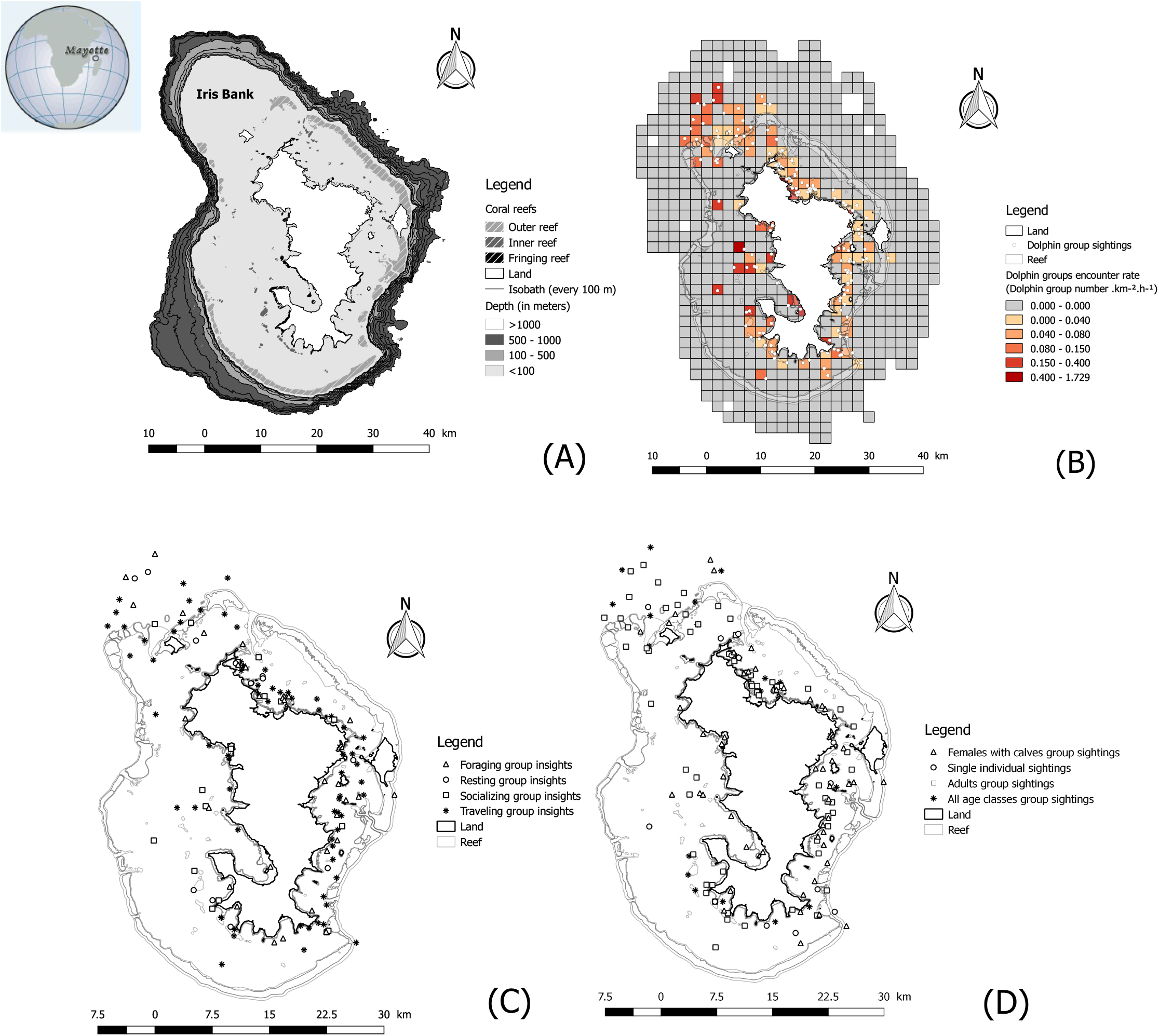
Panel of map detailing the geography of Mayotte and the distribution of dolphin groups, of their behaviors and their structures. **A-** Study area. **B-** Dolphin groups encounter rate. **C-** Behaviors distribution. **D-** Group structures distribution. These maps were produced using QGIS (version 2.18.15).

Field surveys were conducted between 7:00 am and 6:00 pm only on days with good visibility (no rain, Beaufort Sea State < 3; [24]). GPS position was recorded at regular intervals to measure sampling effort. Each time a group was encountered, the time and geographic position were recorded, as well as the behavior and structure of the group.

The behavioral state of a group of dolphins is defined as the activity performed by the majority of the focal group [26]. Activities included: traveling, socializing, foraging, and resting (see [24] for details on sampling). Focal follows were also conducted to investigate behavioral budgets [27]. Focal follows consist of following a group and recording activity every five minutes. Dolphin group structure is defined by the distribution of age and sex within a group. Four age classes can be identified: adults, subadults, calves, and neonates [26]. Group structures include: individuals, adults, females and calves, and groups with all age classes.

### Habitat variables

During the surveys, a number of environmental variables and human activities were collected and used as predictors of dolphin distribution. Variables used included depth, nearest reef type, proxies for prey availability, and measures of human disturbance (boat traffic and dolphin watching activities). These variables were measured and subsequently associated with group sightings using QGIS software version 2.18.15 (including NNJoin and Point sampling tool packages).

Depth data is the main abiotic variable used, it was sampled during the BATHYMAY program [28]. We also measured the distance to the nearest reef as a complementary variable. This variable was highly collinear with bathymetry (r = 0.61) and was only used when bathymetry was not available. Reef types (fringing reef, outer and inner parts of the double barrier reef) were also mapped (see figure 1.A), we associated each observation with its closest reef type.

We also used proxies to estimate prey availability. A first proxy was fish biomass inferred from artisanal fishing data (fisher’s declaration data), this variable will be referred to as “tonnage” in the following analyses. To measure it, the lagoon was divided into activity zones and fishermen were asked about the location of their fishing grounds and the amount of fish caught. For each zone, the weight of fish caught in 2016 was obtained and divided by the area of the zone (values are in kg.km^−2^). Of note, species of the Carangidae family, the main preys of bottlenose dolphin in this area [29], represent a large proportion of fished taxa in the studied areas (21%). To complement this variable, we used the density of fishing sites used by non-commercial fishers. This information was obtained through interviews. Densities (values are in number of fishing spots.km^−2^) were calculated in 4 km^2^ quadrats (see figure 1.B).

Variables measuring human activity were also obtained. First, boat traffic was measured during boat-based surveys conducted between 2014 and 2015. The position of each motorized boat encountered during the transect surveys was recorded. Boat densities were calculated in 16 km^2^ quadrats to avoid autocorrelation bias. These densities were used to calculate encounter rates (values are in number of motorized boats.s^−1^.km^−2^) by dividing the densities by the sampling effort (time spent on each quadrat). Dolphin watching activity was also assessed during focal follow. The number of boats outside and inside a 100 m radius around the dolphin group and the number of swimmers close to the dolphin group were recorded. This 100-meter boundary refers to the current regulation on approaching dolphins in the marine park.

Finally, temporal variations in distribution and habitat preferences were measured. Days were divided into three periods (morning, midday and afternoon) as defined in [24]. Three seasons were distinguished: the rainy season (between June and September), the dry season (between December and March) and the off-season.

### Habitat modeling

Environmental and anthropogenic variables were then used to build models (using RStudio software, version 1.1.383) to explain dolphin distribution and the relationship between occupied habitat and their behavior or group structure. The first model is based on quadrat-scale presence and group count data, while subsequent models are based on sighting data only with neo further spatialization. For the first model, we chose to divide the sampled area into 4 km^2^ quadrats to avoid autocorrelation bias. The number of group sightings and variable values were computed in each quadrat (using its barycenter when distance measures were involved).

A Poisson generalized linear model (GLM) is commonly used to model the distribution of dolphin groups [31]. However, there were presently a large number of null values (no dolphin group present or detected). Therefore, a Hurdle GLM ([32]) was used (package “pscl”). It consists of a combination of a binomial model for presence-absence data and a zero-truncated Poisson model for count data. Using appropriate offsets (quadrat area and sampling effort), this model used the encounter rate as the response variable (number of sightings per unit effort, i.e. time spent surveying per quadrat in our case). The habitat variables used are detailed in table 1. The model fit to the observed data was satisfactory and significantly better than the null model (see table 1).

**Table 1:** Model description and fit to the data. Likelihood ratio tests comparing our models with null models were used to test the significance of fit to the observed data. Such a test does not exist for canonical correspondence analysis, so fit was measured using constraint inertia, which provides a measure of the proportion of variance explained by the variables used. For the other models, McFadden’s pseudo coefficient of determination was used to measure the quality of the fit to observed data. This coefficient, suitable for GLM with a logistic link function, is based on the likelihood ratio. Unlike classical coefficients of determination, it does not vary between 0 and 1; values between 0.2 and 0.4 are usually considered representative of good fit [30]. Levels of significance (p-value): 0 ‘***’ 0.001 ‘**’ 0.01 ‘*’ 0.05 ‘.’ 0.1 ‘ ’ 1

|  | Hurdle GLM | Multinomial GLM | Multinomial GLM | Canonical Correspondence Analysis |
| --- | --- | --- | --- | --- |
| Response variable | Group sightings | Behavioral categories | Group structure categories | Activity budgets |
| Explanatory Variables | Distance to the closest reef,<br>Closest reef type,<br>Tonnage,<br>Fishing spot density | Bathymetry,<br>Closest reef type,<br>Tonnage,<br>Fishing spot density,<br>Motorized boat encounter rate,<br>Daytime,<br>Season,<br>Year | Bathymetry,<br>Closest reef type ,<br>Tonnage,<br>Fishing spot density,<br>Motorized boat encounter rate,<br>Daytime,<br>Season,<br>Year | Bathymetry,<br>Tonnage,<br>Fishing spot density,<br>Motorized boat encounter rate,<br>Boats number within 100 meters,<br>Boats number over 100 meters,<br>Swimmers number,<br>Daytime,<br>Season,<br>Year |
| Likelihood-ratio test | $<10^{-15}$ | 0.0248 | 0.0316 | - |
| Pseudo R-squared<br>(or constraint inertia) | 0.197 *** | 0.195 * | 0.157 * | 0.507 |

To determine whether habitat variables affected the observed behaviors or group structures, we used multinomial GLMs ([33], [34]) using the “nnet” package. The different behavioral categories or group structures were considered as response variables. The habitat variables used are detailed in table 1. The model fit to the observed data was satisfactory and significantly better than the null model (see table 1).

An analysis of focal follows was also used to understand which habitat variables influence activity budgets (e.g., [35]). For each focal follow, the mean values of the habitat variables were calculated over the time series. The relationship between the activity budget and averaged habitat variables was then explored by canonical correspondence analysis using the “vegan” R package. The habitat variables used are detailed in table 1. The model fit to the observed data was satisfactory (table 1).

To analyze the significance of the variables’ effects, we used analyses of deviance (R package “car”) for GLMs, that use likelihood ratio tests between models with and without the tested variables. For canonical correspondence analysis, a permutation F-test was used (R package RVAide-Memoire). For qualitative variables, post hoc tests were used to determine which class differed from each other: we used a t-test on pairwise contrasts for Hurdle GLMs (R package emmeans) and pair-wise F-tests for canonical correspondence analysis. The effects of quantitative and qualitative variables were further analyzed for multinomial GLMs: we used odd ratios to compare effect sizes (ratio of the observation probability between two categories of the response variable when the explanatory variable of interest varied by one unit for a quantitative variable or from one category to another for a qualitative variable), the significance of odd ratios was tested by z-tests.

## Results

### Sampling effort and data set

The area surveyed was approximately 2,740 km^2^. A total of 1940 hours were spent surveying the Mayotte lagoon and adjacent waters. From 2004-2009 and 2015-2016, 160 groups of Indo-Pacific bottlenose dolphins were detected (figure 1.B). During encounters, dolphins were traveling (n=72, 45%), foraging (n=31, 19.4%), socializing (n=17, 10.6%), and resting (n=12, 7.5%, figure 1.C). Behaviors of some groups could not be identified (n=28, 17.5%). Group composition was variable and included groups of adults only (n=69, 43.1%), groups of females with calves (n=50, 31.3%), groups of all ages (n=21, 13.1%), and single individuals (all adults, n=13, 8.1%, figure 1.D). Group structures could not be identified for some sightings (n=7, 4.4%). Most sightings are distributed inside the lagoon or on the Iris Bank at a mean depth of 35 m (figure 1.B).

### Species distribution model

To estimate the effects of habitat variables on the encounter rate of dolphin groups, we implemented a species distribution model (see Materials & Methods). Analysis of deviance shows that tonnage (a proxy for prey abundance), reef type, and distance to reefs all have a significant effect on the probability of dolphin encounters. This probability increases significantly as distance to the nearest reef decreases (see figure 2.A) and is significantly higher near inner and fringing reefs (see figure 2.B). It also increases with increasing tonnage (see figure 2.C). There is no significant effect or trend in the density of the fishing area on the probability of presence. Distance to the nearest reef is the only variable that affects dolphin group density: dolphin group density increases significantly as distance to the reef increases (see figure 2.D). There are no significant effects or trends for the other three variables. These effects show that bottlenose dolphins are mainly found within the lagoon (except for Iris Bank), in shallow environments, and preferentially in relative proximity to coral reefs, but not directly upon them (see figures 1.A and 1.B).

**Figure 2:**
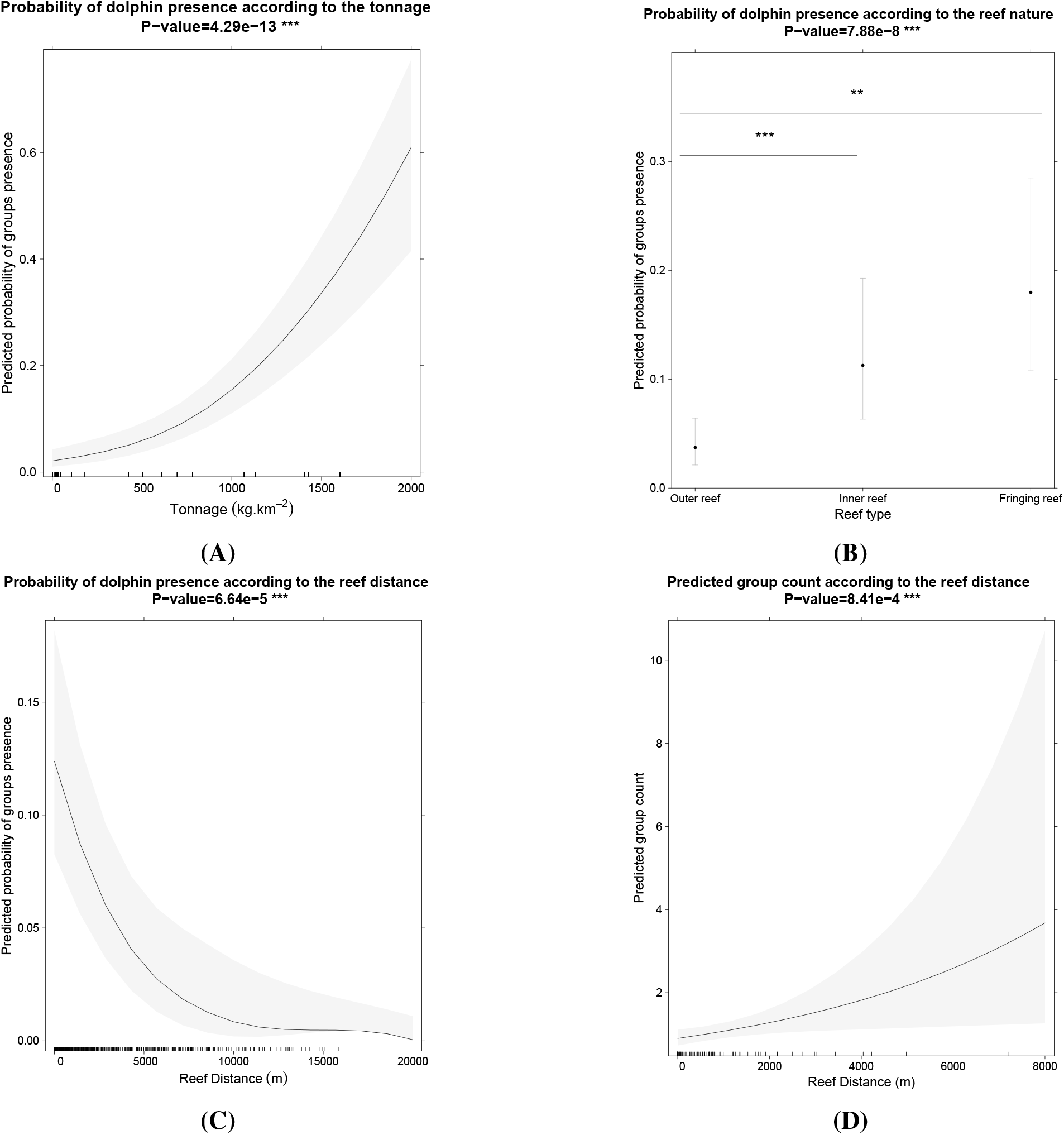
Panel of graphs showing the effect of habitat variables on dolphin group presence and density, obtained from the Hurdle GLM. Only the plots of variables with significant effects are shown here. Graphs A to C were obtained from the binomial component of the hurdle GLM, testing the effect of habitat variables on the probability of group presence. Graph D was obtained from the binomial component of the hurdle GLM, testing the effect of habitat variables on group density in cases where at least one group was observed. The shaded area and gray bars represent the 95% confidence interval computed from a pointwise method. The black bars on the x-axis represent the distribution of the habitat variables. P-values obtained from the analysis of deviance are given for each variable above the graphs. For qualitative variables with significant effects, we have provided pairwise test levels of significance, above black horizontal bars bridging the categories being compared. These plots were generated using the “effects” package. Levels of significance (p-value): 0 ‘***’ 0.001 ‘**’ 0.01 ‘*’ 0.05 ‘.’ 0.1 ‘ ’ 1

### Habitat Variables Impact on Behaviors

We then sought to understand the relationship between habitat variables and the behaviors of the observed dolphin groups. The first approach used was to consider the distribution of all behaviors as a function of habitat variables using a multinomial model (see Materials & Methods). Analysis of deviance shows a significant effect of time of day and nearest reef type (see figures 3.A, 3.B). Analysis of significant odd ratios shows an increase in the proportion of socializing groups in the afternoon compared to the morning, an increase in the proportion of foraging groups in the midday compared to the afternoon, and a decrease in the presence of resting groups in the midday compared to the rest of the day (see figure 3.A). This distribution of behaviors throughout the day reflects a daily cycle of activities. There is also a significant effect of reef type on dolphin occurrence probabilities. Socializing groups had higher probabilities of occurrence near inner reefs compared to fringing reefs. Resting groups had higher probabilities of occurrence near outer reefs compared to inner reefs. Foraging groups had higher probabilities of occurrence near the fringing reef compared to other reefs (see figure 3.B, and figures 1.A, 1.C). The other variables did not have a significant effect on dolphin abundance, including the encounter rate of motorized boats. These effects support the existence of spatial and temporal segregation of behaviors, with different functional habitats being identified relative to the nature of the closest reef.

**Figure 3:**
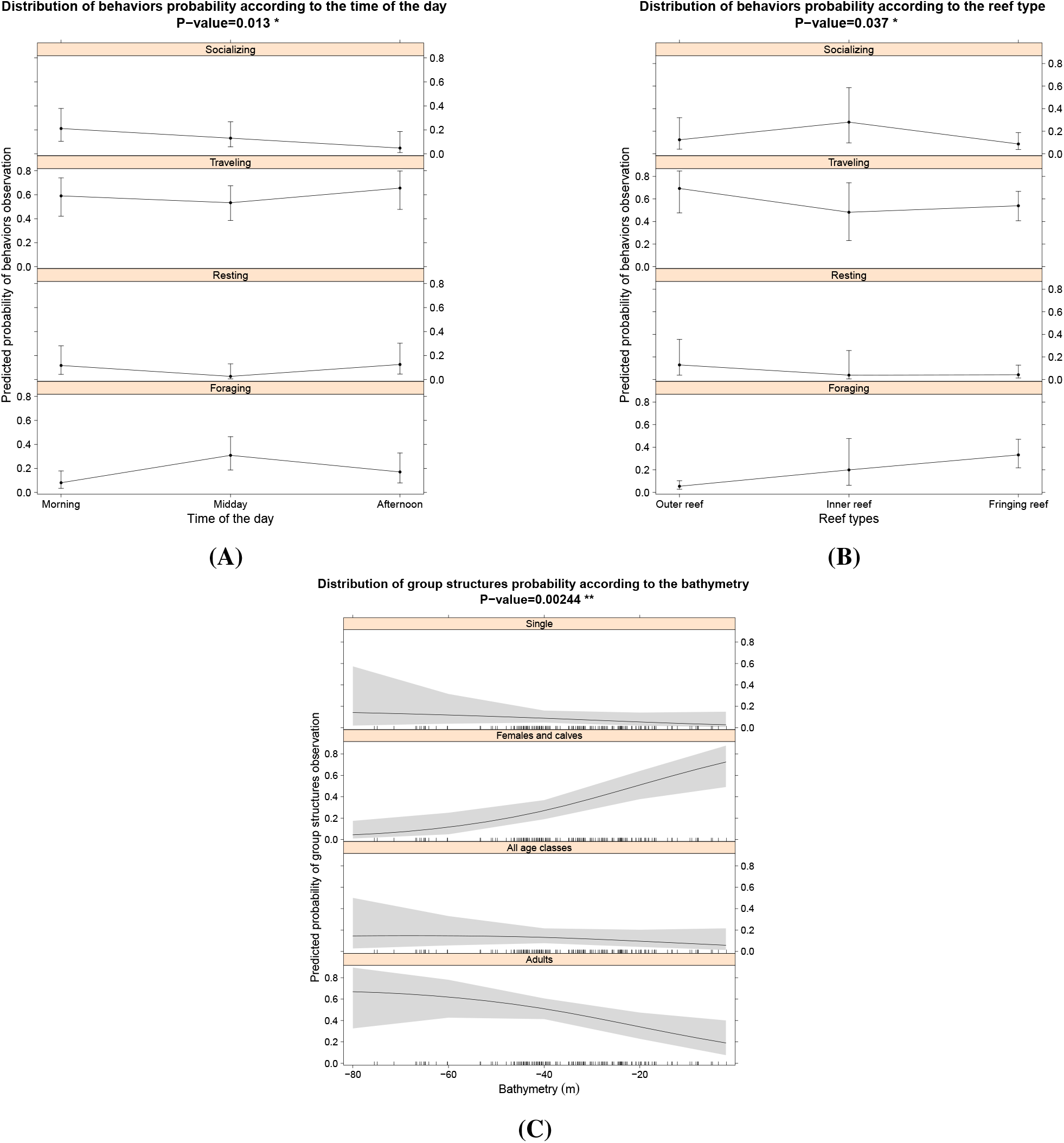
Panel of graphs showing the effect of habitat variables on dolphin group behavior and composition, obtained from a multinomial GLM. Only the plots showing significant effects are shown here. Plots A and B refer to effects on behaviors, and plot C refers to effects on group structure. The shaded area and gray bars represent the 95% confidence interval, computed from a pointwise method. The black bars on the x-axis represent the distribution of the habitat variables. P-values obtained from the analysis of deviance are given for each variable above the graphs. These plots were generated using the “effects” package. Levels of significance (p-value): 0 ‘***’ 0.001 ‘**’ 0.01 ‘*’ 0.05 ‘.’ 0.1 ‘ ’ 1

Focal follows were also used to identify the potential effects of habitat variables on behavior (see Materials & Methods). We used a canonical correspondence analysis, which shows significant effects of fishing site density and time of day on activity budgets. There is a significantly higher proportion of foraging behaviors and a lower proportion of traveling behaviors for focal follows with high fishing spot density values (a proxy for prey abundance) observed along the track. According to post hoc tests, the effect of the time of day is mainly focused on the comparison between the afternoon and the rest of the day, with a significant increase in the proportion of socializing behavior in focal follows that occurs during the afternoon, in concordance with what was observed for the multinomial GLM.

### Habitat Variables Impact on Group Structures

As with behavior, we used a multinomial model to determine which habitat variables influence group structure (see Materials & Methods). Analysis of deviance shows a significant effect of depth on the probability of occurrence of females with calves: the encounter probability of these groups tends to be higher at low depths (see figure 3.C, and figures 1.A, 1.D). The other variables have no significant effect on the group structure.

## Discussion

Our analyses suggest that the studied population has a strong preference for coastal waters near coral reefs, in shallow waters. This confirms what was found in a previous study [24]. Inshore habitats should be associated with high prey availability given the feeding habits of bottlenose dolphins [29], which is confirmed by a preferential distribution of dolphin groups in areas with high fishing activity (i.e. high fish productivity). Behaviors seem to follow a daily cycle, which has been observed in other populations as well, but not systematically in the same order [36]. Behaviors also seem to have distinct spatial distributions, being preferentially located near different reef types. Habitat is also related to group composition: groups with females and calves prefer shallow habitats. This distribution can be better understood by the theory of optimal foraging [11]. The risk of predation would be rather low in this population [37], so according to the ideal free distribution [12] the main distribution drivers should be prey availability and competitive pressure. As expected, groups seem to be located in areas of high prey productivity (as indicated by the tonnage values). Interspecific competition with other dolphins is absent because of ecological niche partitioning around Mayotte [24] and intraspecific competition is probably reduced by habitat partitioning [18]. To obtain more precise information on the importance of prey distribution, variables directly related to prey abundance should be used instead of information based on fishing activity, but this would require specific surveys to be conducted for future studies. It would be interesting to also test the accessibility of prey between reefs using rugosity metrics (which measures the availability of shelters for prey, [38]). These results highlight that prey depletion due to overfishing [19] around Mayotte may become a major concern, so it would be interesting in the future to study the impact of the temporal evolution of prey availability on dolphin distribution.

We cannot conclude on the validity of the risk-disturbance hypothesis [14] for the present population as we did not observe a significant effect of boat traffic on the distribution and abundance of dolphin groups. This may be due to the poor spatio-temporal resolution of the variable. However, the avoidance of disturbance could also be temporal [16]: some investigations on temporal influences could be carried out to complement this study. However, the absence of negative effects could also reflect that there is no other option than to stay in this habitat for dolphins: the study of the potential physiological effects of anthropogenic disturbance [39] could then be valuable, with further consideration of complementary variables on human disturbance as acoustic measures of noise pollution [40]. In addition, to better understand the impact of human activities on dolphin distribution and abundance, complementary investigations on habitat degradation status (using proxies such as coral, reef, seagrass, and mangrove cover) could be conducted in future studies given the recent pressures on [21].

In this population, habitat selection also seems to depend on group structure and activities: different functional habitats [12] seem to be used. Of note, we must be cautious about some of these effects, as sample sizes for some behavioral categories (resting) or group types (single individuals) were relatively small, limiting our statistical power. We observed that females with calves preferred nursery habitats at lower depths, which presumably provide them with better protection from male aggression, predation risk, and possibly boat traffic [41]. They also prefer environments with high fish productivity more than other group structures, probably because they have high resource requirements and low energy reserves [42]. Other functional habitats dedicated to different behavioral activities appear here, as the distribution of resting, socializing, and foraging behaviors are distributed around different coral reef types. In particular, foraging habitats appear to be distributed in coastal environments with high fish productivity, whereas socializing and resting occur further from shore, near the fringing and outer reef, respectively. The observed effect of fishing site density on the activity budget (increasing proportion of foraging behavior relative to traveling behavior when the density of fishing spots increased) suggests that foraging behavior is indeed more likely to occur in areas of high prey density. The effect on the proportion of traveling behavior could be explained by considering this behavior as a transitional behavior, and in particular as a way to move from one foraging site to another. Other variables, such as human disturbance, did not appear to affect activity budgets here, but further analyses of these budgets, such as behavioral transition models based on Markov chain analyses [43], could be useful to better understand the impact of these disturbances.

## Author contribution & Acknowledgments

MB performed all statistical analyses and write the manuscript. JJK designed the study, collected data collected from 2004 to 2008, reviewed the manuscript, and supervised analyses and manuscript writing. JW collected data from 2014 to 2015, provided environmental data, and contributed to the design of the study. SJ contributed to the design of the study. All authors reviewed the manuscript.

